# Single-nucleus transcriptomics resolves differentiation dynamics between shoot stem cells and primary stem

**DOI:** 10.1101/2024.08.06.606781

**Authors:** Sebastián R. Moreno, Martin O. Lenz, Elliot M Meyerowitz, James CW Locke, Henrik Jönsson

## Abstract

The shoot apical meristem (SAM), located at the plant apex, is accountable for the formation of above-ground organs such as leaves, stem and flowers. Although transcriptional profiling has elucidated some cell-types observed within stems or flowers, the differentiation transcriptional dynamics from shoot stem cells to multiple cell identities remain unknown. We employed a single-nucleus RNA-sequencing approach to assess the transcriptional heterogeneity and cell differentiation processes within the SAM. By collecting dissected inflorescence meristems, we constructed an inflorescence single-nucleus SAM atlas from *Arabidopsis thaliana*. Our analysis unveiled regulatory elements for most previously known cell types such as the boundary domain, vasculature, early primordia, epidermis and internal stem cells. We also identified previously unobserved transcriptional profiles, revealing that the stem cortex is defined early within forming primordia. Moreover, trajectory inference analysis allowed us to capture spatial control of S-phase machinery by floral homeotic genes and differentiation gene expression dynamics from internal shoot stem cells toward internal layers such as cortex, cambium, xylem and phloem. The results advance our understanding of the cellular and transcriptional heterogeneity underlying the cell-fate transcriptional dynamics shaping shoot organs and architecture.

## INTRODUCTION

In contrast to many animals, whose organogenesis is confined to embryonic or larval stages, plants generate organs throughout their lifespan. This continuous organogenesis is orchestrated by stem cells situated within specialized regions known as meristems (Nägeli et al., 1858; Newman, 1965). The shoot apical meristem (SAM), located at the plant apex, is accountable for the formation of above-ground organs such as leaves, stem and flowers (Traas and Vernoux, 2002).

In angiosperms, SAM stem cells reside in two but overlapping zones: the Central Zone (CZ) and the Organizing Centre (OC), the first apical and the second only in internal layers (Laux et al., 1996; Reddy and Meyerowitz, 2005). These stem cell reservoirs displace daughter cells outwardly toward the periphery and internal stem, where they will differentiate into the various cell types observed in aerial organs (Burian et al., 2016). Most CZ lineages within the SAM are characterized by anticlinal division, ensuring isolated cell lineages within the epidermal layer and subepidermal layers (Meyerowitz, 1997; Reddy et al., 2004). Conversely, OC lineages undergo both periclinal and anticlinal divisions (Bencivenga et al., 2016), forming the diverse compendium of cell types present in inner layers of primary stem such as cortex, pith and vascular bundles (Reviewed in Miyashima et al., 2012). The mechanisms and transcriptional dynamics linking shoot stem cells to the diverse cell types observed in the primary stem have remained largely unknown.

In *Arabidopsis thaliana,* the CZ is positioned centrally in the SAM and is demarcated by the expression of the *CLAVATA 3* (*CLV3*) peptide-encoding gene. *CLV3* is expressed in the epidermis, and also in internal layers where it overlaps with cells expressing the RNA for the transcription factor WUSCHEL (WUS), which defines the OC domain (Mayer et al., 1998; Fletcher et al., 1999; Schoof et al., 2000). Beyond these stem cell pools, other marker genes and/or characteristic morphological domains demarcate various cell types in the SAM, such as the Boundary Domain (BD) in the boundary between new floral primordia and the SAM (Aida et al., 1997), an auxin-responsive domain marking primordial (leaf or floral) initiation (Snow and Snow, 1937; Benková et al., 2003), a peripheral zone (PZ) surrounding the OC (Meyerowitz 1997; Tian et al., 2019) and a rib zone beneath the OC that gives rise to the stem pith (Bencivenga et al., 2016). Furthermore, some vascular genes expressed in the stem are also expressed in specific domains within the SAM, defining pro-cambial strands during primordial formation (Mor et al., 2022; Sanchez et al., 2012; L. Zhang et al., 2014), or exhibiting an internalized string pattern starting from the rib zone (Wahl et al., 2010). Although some cell types within the stem and flowers such as procambium have been identified to originate within the SAM, whether there are other cell-types defined within the meristematic domains is unknown.

In recent years, single-cell RNA-sequencing approaches have been employed to characterize the cellular heterogeneity within the shoot apex across diverse plant species such as Arabidopsis (Zhang et al., 2021; Xu et al., 2024), rice (Zong et al., 2022), Populus (Conde et al., 2022; Du et al., 2023) and maize (Satterlee et al., 2020; Xu et al., 2021). While these studies have provided invaluable insights into the diversity of cell types within the shoot apex, the identification of shoot stem cells has been elusive. In an effort to capture more cells expressing the key stem cell markers CLV3 and/or WUS, some studies have utilized mutant backgrounds characterized by over-proliferation in the inflorescence meristem (Neumann et al., 2022; Xu et al., 2024). However, the use of mutants with markedly different morphological features and patterns of gene expression compared to wild-type would be expected to hamper the comprehensive reconstruction of differentiation trajectories originating from wild-type shoot stem cells.

In this study, we employed a single-nucleus RNA-sequencing (snRNA-seq) approach from finely dissected wild-type inflorescence meristems to unravel the transcriptional heterogeneity and cell differentiation processes within the SAM. Our analysis unveiled regulatory elements for most previously known cell types such as boundary domain, vasculature, early primordia, epidermis, S- and G2/M-phase and internal stem cells. In addition, we observed that cortex is also defined within the SAM, along with vasculature during primordium formation. Furthermore, trajectory analysis allowed us to reconstruct cell-cycle gene expression dynamics in the SAM and allowed us to reconstruct the transcriptional landscape from shoot stem cells to diverse differentiated cell identities such as cortex, xylem, phloem, cambium and early primordia. This advances our understanding of the cellular and transcriptional correlates of cell fate acquisition from shoot apical meristem stem cells.

## RESULTS

### Diverse biological functions are shaped by gene expression heterogeneity in the shoot apical meristem

To explore the cellular diversity within the shoot apical meristem (SAM) of *Arabidopsis thaliana*, we employed a snRNA-seq approach using dissected wild type meristems. The dissection process involved excising all flowers beyond stage 4 (Smyth et al., 1990; Alvarez-Buylla et al., 2010) and sectioning the SAM around 150-200 μm below the apex of the meristematic tissue (Figure 1A). Immediately following dissection, tissues were cryopreserved in liquid nitrogen for subsequent nuclear isolation (Methods). The isolated nuclei were then processed using the 10X Genomics Chromium platform and subsequently sequenced using the Illumina NovaSeq 6000 system. Three independent batches were sequenced (Supplementary Figure 1A and 1B). Following data integration and quality control, a total of 7,295 single-nucleus transcriptomes covering 20,454 genes was obtained. Unbiased clustering analysis of transcriptional heterogeneity led to the classification of nuclei into 16 cell type clusters (Figure 1B, Methods, Supplementary Table 1). Increasing clustering resolution diminished the specificity of cell-marker assignment (Supplementary Figure 1C).

**Figure 1.**
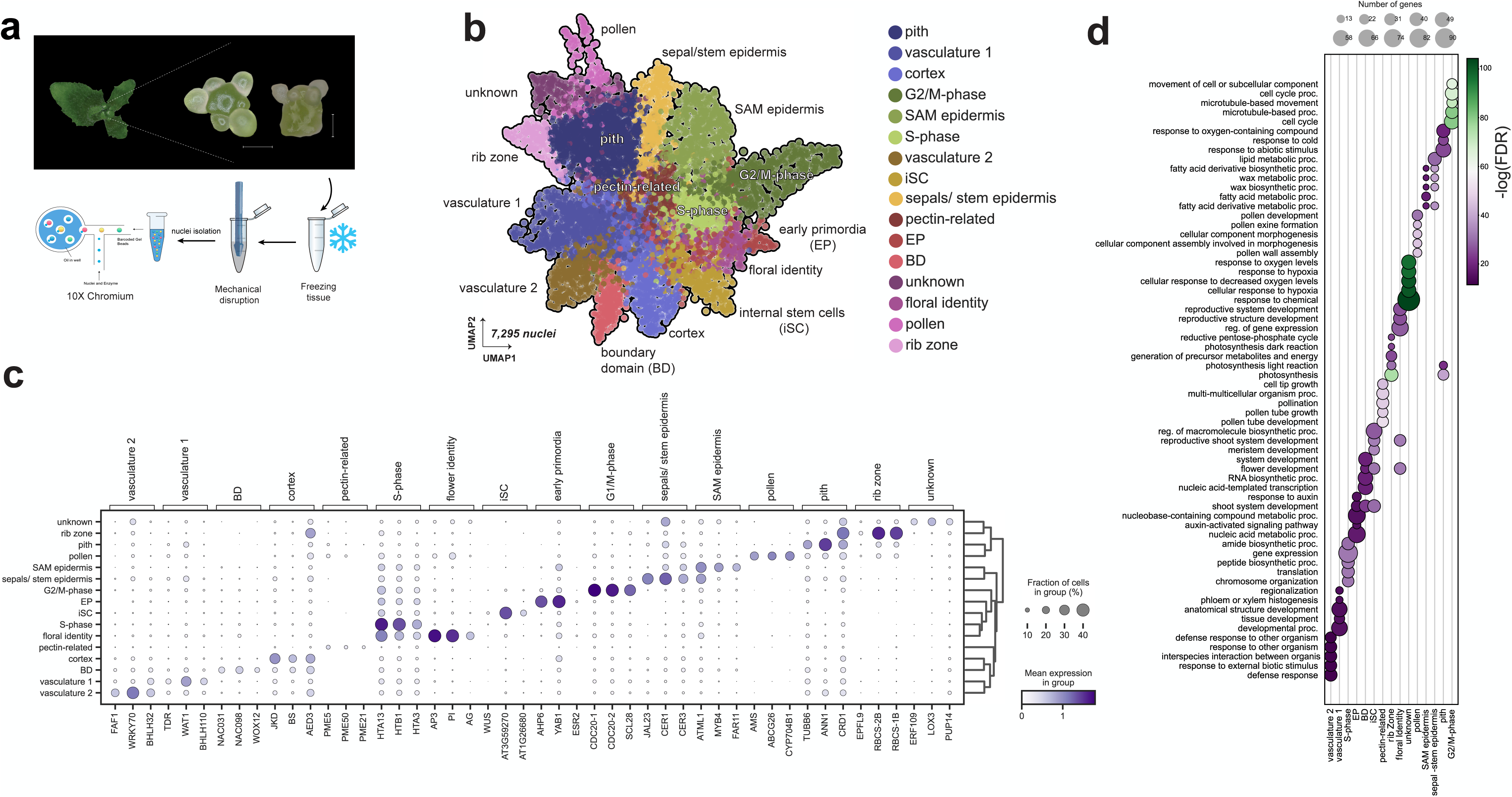
snRNA-seq resolves cellular heterogeneity within the *Arabidopsis thaliana* shoot apical meristem. a) Schema with general workflow used for single-cell transcriptome analysis from dissected *Arabidopsis thaliana* SAMs. Floral meristems with distinct sepals were removed (> stage 4). SAMs were frozen and then mechanically disrupted by pestle homogenization. Nuclei were FACS sorted before loading onto 10X Chromium chips. Scalebar = 100 µm. b) Uniform manifold approximation (UMAP) and projection of cellular heterogeneity in SAM tissue (7,295 cell meristematic nuclei). c) Dot plot showing selected marker genes for each cell cluster. The colour bar indicates the gene-normalized gene expression. Dot size indicates the fraction of cells expressing marker genes. d) GO enrichment analysis from the top 100 marker genes per cluster. 5 top GO terms sorted by Fold Enrichment were selected per cluster.

We used prior knowledge of where certain identified marker genes are expressed within the SAM to label the different gene expression clusters (Figure 1C, Supplementary Figure 2). The BD cluster was identified by the expression of *CUP-SHAPED COTYLEDON 2* and *3* (*CUC2* and *CUC3*) (Takeda et al., 2011). Another cluster that exhibited expression of YABBY1 (*YAB1)*(Goldshmidt et al., 2008)*, HISTIDINE PHOSPHOTRANSFER PROTEIN 6* (*AHP6*) (Besnard et al., 2014), *DORNROESCHEN-LIKE* (*DRNL*)(Dai et al., 2023) and numerous other genes associated with primordium formation was designated as Early Primordia (EP) (Figures 1C and 3A). Although a discrete cluster for epidermal stem cells located in the CZ was not identified, we distinguished between meristematic and differentiated epidermis (Figure 1C). While epidermal marker genes such as *MERISTEM LAYER 1* (*ATML1)* and *PROTODERMAL FACTOR 1* (*PDF1*) were expressed in both clusters (Figure 1C, Supplementary Figure 2A and 2B) (Sessions et al., 1999; Fal et al., 2021), specific marker genes for each epidermal cluster were detected. We named Sepals/stem Epidermis a cluster with cells expressing differentiated epidermal cell marker genes such as *ECERIFERIUM 1* and *3* (*CER1* and *CER3*) and other genes related to cuticular wax biosynthesis (Supplementary Figure 2A) (Bourdenx et al., 2011; Pascal et al., 2019; Vadde et al., 2024). In a second epidermal cluster, cells expressing *CLV3* and *KANADI 1* (*KAN*) were observed, all of which have been shown to be expressed in the SAM epidermis (Supplementary Figure 2B) (Caggiano et al., 2017). Rib zone cells were identified by the expression of *EPIDERMAL PATTERNING FACTOR-LIKE 9* (*EPFL9*), which has been observed to be expressed underneath the SAM (Kosentka et al., 2019). The Pith cluster was identified using available spatial transcriptomic data that defined pith shoot apex (Du et al., 2023). The pith shoot apex cluster identified by spatial transcriptomics overlaps with marker genes observed in our dataset (Supplementary Figure 2D and 3). Cortex cells were defined by the expression of *JACKDAW* (*JKD*)*, ASPARTYL PROTEASE AED3* (*AT1G09750*), and *CELL WALL/VACUOLAR INHIBITOR OF FRUCTOSIDASE* (*c/VIF2*), among other genes previously observed to be expressed in root cortex (Figures 1C and 4A) (Hassan et al., 2010; Shahan et al., 2022; Nolan et al., 2023). The Vasculature 1 cluster was defined by the expression of SAM procambium gene *HOMEOBOX GENE 8 (ATHB8)* (Kang et al., 2003; Sanchez et al., 2012) and genes previously identified in root vasculature such as CALLOSE SYNTHASE 8 (CALS8) (Ross-Elliott et al., 2017) and *PERICYCLE FACTOR TYPE-A5* (*PFA5*) (Figures 1C and 2A) (Zhang et al., 2021). The Vasculature 2 cluster is related to other SAM- and stem-related vasculature genes such as *FANTASTIC FOUR 1 and 3* (*FAF1*, *FAF3*) and *XYLOGLUCAN ENDO-TRANSGLYCOSYLASE/HYDROLASE 4* (*XTH4*) (Supplementary Figure 2E) (Wahl et al., 2010; Kushwah et al., 2020). We also observed a cluster of cells with the expression of several genes related to pectins such as *PECTIN METHYLESTERASE 21* (*PME21*), *PME50*, *PME13*, *PME4*, *PME5*, *RALFL19* and *RALFL4* (Supplementary Figure 2F). *PME5* expression has been previously reported in the SAM with a scattered pattern (Peaucelle et al., 2011). We named this cluster the Pectin-related cluster. A group of cells differentially expressing floral homeotic genes such as *APETALA 3* (*AP3*), *PISTILLATA* (*PI*), *AGAMOUS* (*AG*) and *SEPALLATA 1-3* (*SEP1-3*) were observed. Floral homeotic genes have distinct expression patterns in different domains of flower primordia (Jack et al., 1994; Krizek and Meyerowitz, 1996; Liu et al., 2009; Urbanus et al., 2009). We named this group of cells the Floral Identity cluster (Figure 1B-C and Supplementary Figure 2G). There was a distinct cluster of cells that showed co-expression of *WUS*, *AT3G59270* and *AT1G26680* (Figure 1B-C), resembling the known OC domain (Figure 1D) (Supplementary Figure 2H) (Ung et al., 2011). According to the expression of *AT3G59270*, this cluster is spatially located in the OC but extends a few cells more broadly than the WUS domain. We designated this cluster as internal Stem Cells (iSC). S-phase and G2/M-phase clusters were identified by the expression of histone-related genes and mitotic marker genes, respectively (Supplementary Figure 2I and 2J). One cluster, Unknown, remains associated with any known domain due to the lack of available reporter lines (Supplementary Figure 2K) and another cluster consists of cells that express pollen-related genes, perhaps due to the occasional pollen grain found at the top of the meristem post-dissection (Supplementary Figure 2I).

**Figure 2.**
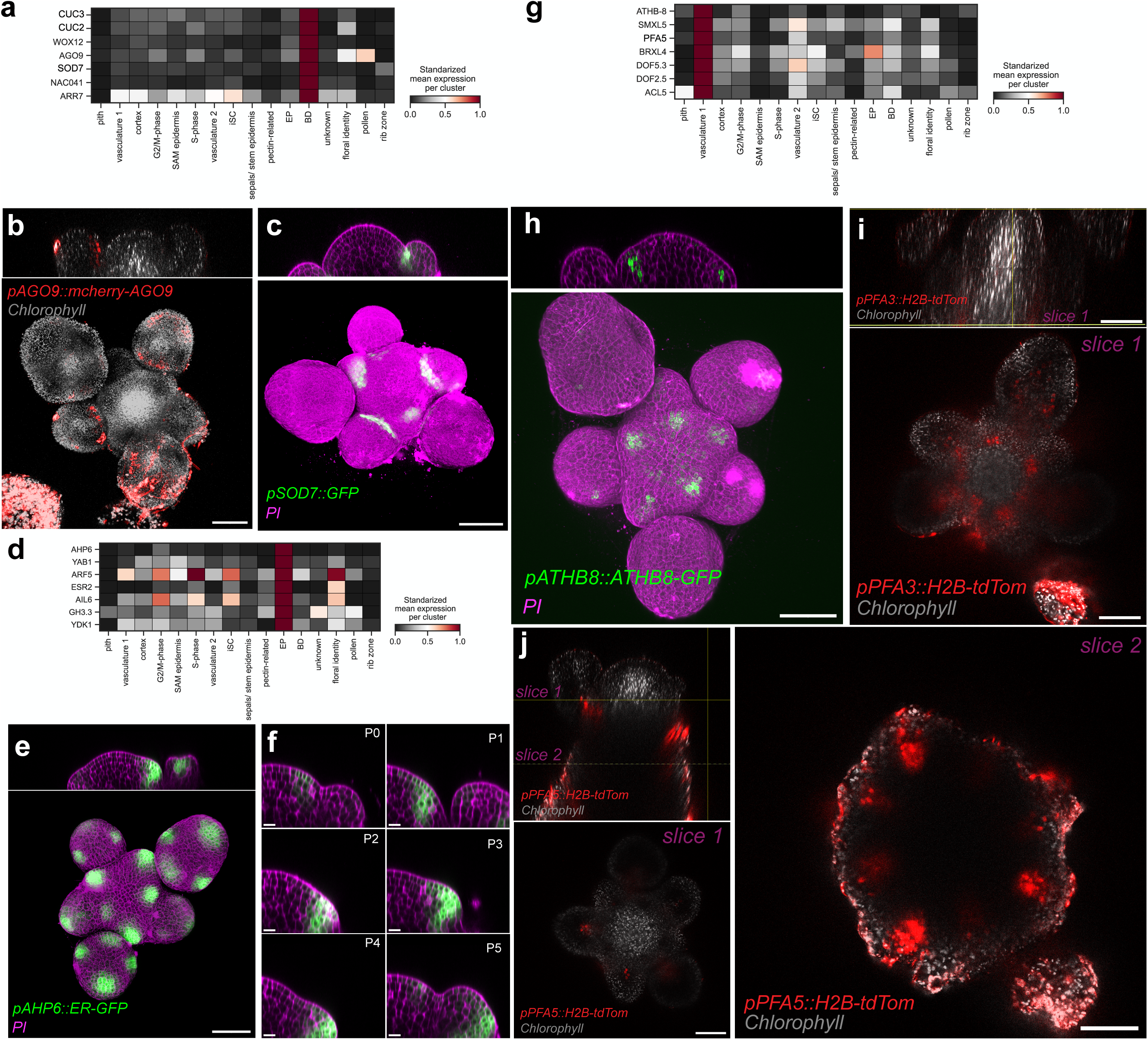
Cell clusters captured cellular identities at different differentiation states. a) Heatmap plot illustrating selected marker genes from the BD cluster. b) Orthogonal projection (top) and maximal projection (bottom) of fluorescence from the *pAGO9::mCherry-AGO9* reporter in the SAM. c) Orthogonal projection (top) and maximal projection (bottom) of the pSOD7::GFP reporter line in the SAM. d) Heatmap plot illustrating selected marker genes from the EP cluster. Each gene is normalized by its own expression. e) Orthogonal projection (top) and maximal projection (bottom) of fluorescence from the *pAHP6::ER-GFP* reporter in the SAM. f) Orthogonal primordia slices of *pAHP6::ER-GFP* reporter line following primordium formation from incipient primordia to primordium number 5: scale bars, 5 µm. g) Heatmap plot illustrating selected marker genes from the Vasculature 1 cluster. h) Orthogonal projection (top) and maximal projection (bottom) of *pATHB8::ATHB8-GFP* reporter line in the SAM. i) Orthogonal projection of *pPFA3::H2B-dTomato* reporter line in the SAM with a confocal slice labelled by a yellow line. j) Orthogonal projection of *pPFA5::H2B-tdTom* reporter line in the SAM with two confocal slices at different stem depths (bottom and right panels). scale bars, 50 µm.

**Figure 3.**
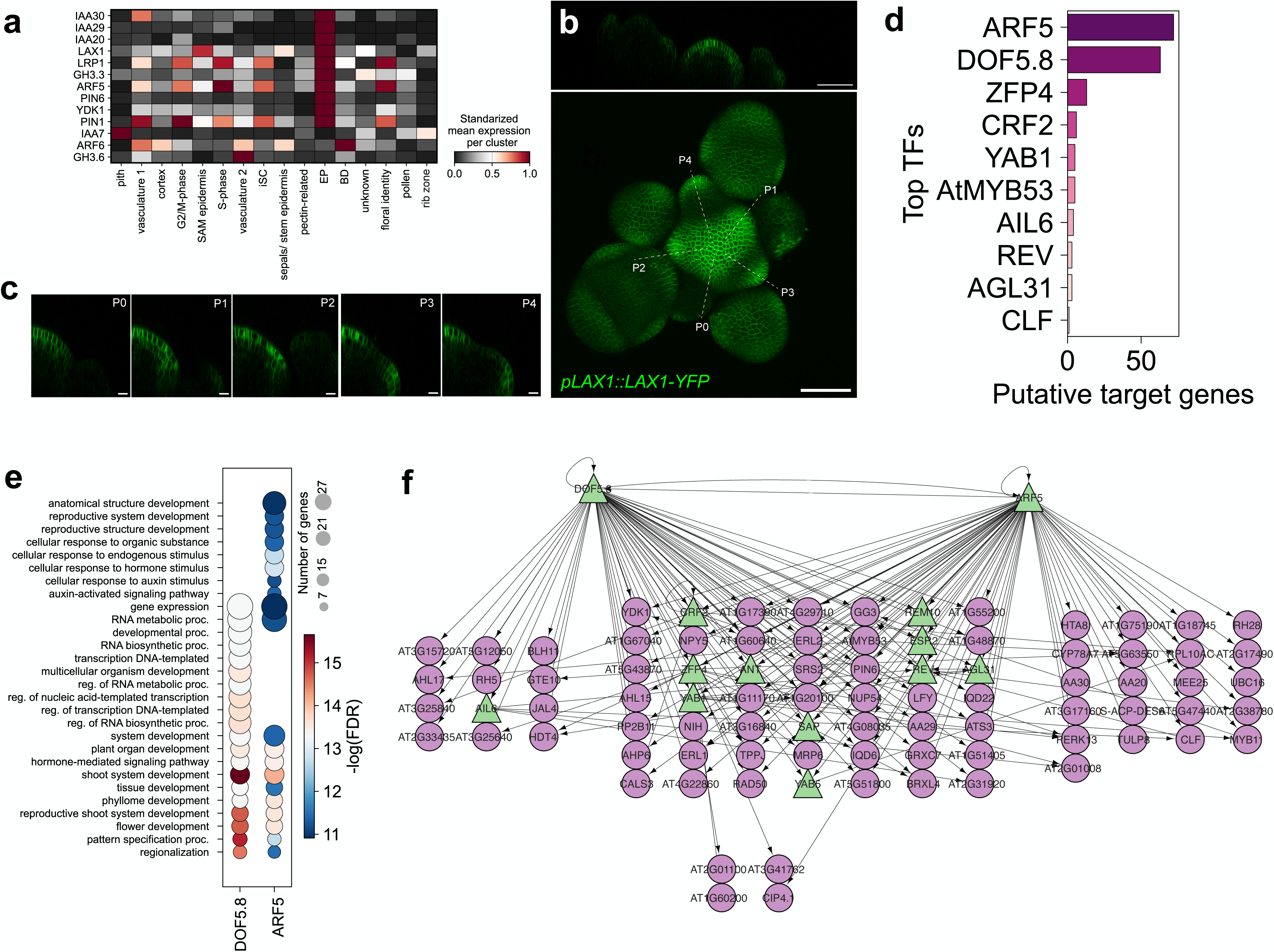
ARF5 and DOF5.8 as the main regulators of primordium formation in the SAM. a) Heatmap plot illustrating normalized expression values of auxin-related marker genes. b) Orthogonal projection (top) and maximal projection (bottom) of *pLAX1::LAX1-YFP* reporter line in the SAM. c) *pLAX1::LAX1-YFP* reporter line following primordium initiation ordered from incipient primordia. Scale bars, 5 µm. d) Ranking of TF marker genes within EP cluster based on the number of putative target genes. e) GO analysis for ARF5/MP and DOF5.8 target genes. f) GRN of EP cluster from top 100 marker genes for EP cluster. TFs are showed with green arrows. Size of the triangle is proportional to the number of target genes.

**Figure 4.**
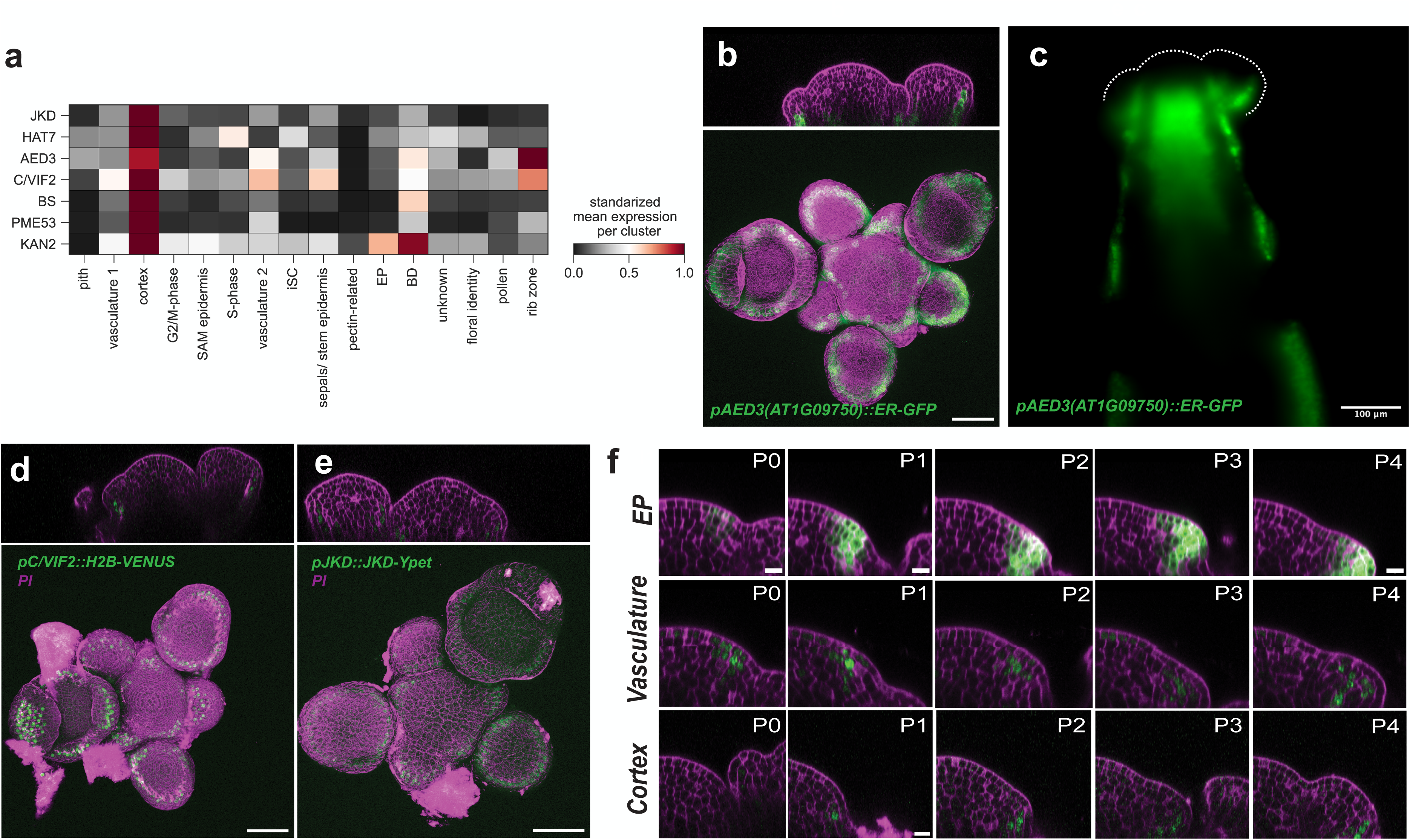
Cortex and vasculature identities diverge during primordium formation within the SAM. a) Heatmap plot illustrating selected cortex-related marker genes with normalized gene expression by gene. b) Maximal projection of SAM from *pAED3(pAT1G09750)::ER-GFP* reporter lines stained with propidium iodide (PI). c) Orthogonal projection of light-sheet image of *pAED3(pAT1G09750)::ER-GFP GFP* reporter lines. d) Orthogonal and maximal projection of SAM from pC/VIF::H2B-VENUS reporter lines stained with PI. e) Orthogonal and maximal projection of SAM from pJKD::JKD-Ypet reporter lines stained with PI. f) Primordium formation in the *pAHP6::ER-GFP* reporter line (upper panel), *pATHB8::ATHB8-GFP* reporter line (middle panel) and *pJKD::JKD-Ypet* reporter line (bottom panel) following the phyllotactic pattern. scale bars for primordium initiation, 5 µm. Scale bars, 50 µm

We then asked whether there are biological functions significantly enriched in the GO categorizations of the marker genes identified in each cell type. Different biological functions were significantly enriched per cluster with a low level of overlap (Figure 1D). Most of the observed GO terms are associated with the cell identities previously assigned based on unbiased clustering and marker gene analysis. For example, iSC marker genes were enriched in meristem development reproductive shoot system development and many other GO terms related to the stem cell niche. The EP cluster was enriched in genes related to auxin-activated signalling pathways and response to auxin terms. The Vasculature 1 cluster showed differentially enriched GO terms such as phloem or xylem histogenesis as expected. Thus, GO term enrichment analysis accurately separated clusters by biological function, validating the differential expression analysis that identified the marker genes in each cluster.

### Cell identities are discernible at different stages of differentiation

To assess the accuracy of unbiased clustering, we examined the expression pattern of predicted differentially expressed (DE) marker genes. In the BD cluster, the gene *SUPPRESSOR OF DA1-1 (SOD7/NGAL2)* has been previously characterized as repressing CUC expression during the initiation of axillary meristems (AMs) and it was also observed as a marker gene in our approach (Nicolas et al., 2022) (Figure 2A and 2C). Another differentially expressed gene in BD cluster was *ARGONAUTE 9* (*AGO9*). Using a translational reporter line, we observed *AGO9* expression in the boundary between sepals and the floral meristem in developing flowers (Figure 2B), suggesting similar boundary domain identities at various stages of flower development. Previous work has shown that the boundary domain between SAM and FM share a similar expression pattern with sepal boundaries (Refahi et al., 2021), supporting the observation that cells from both domains are found in the same cluster.

Within the EP cluster, we observed the expression of previously characterized genes such as *AHP6*, *YAB1* and other genes associated with primordium formation (Figure 2D). Although *YAB1* has been predominantly associated with the abaxial patterning in primordium and leaf (Goldshmidt et al., 2008), the spatial gene expression pattern of *AHP6* indicates that this cluster captured nuclei from the entire early primordium, without distinguishing between abaxial or adaxial sides (Figure 2E and 2F).

The Vasculature 1 cluster showed expression of genes previously described as active during vascular differentiation in roots and shoot (Figure 2G). *ATHB8* RNA has been observed in the procambial strand within the SAM connecting floral primordia with primary stem (Kang et al., 2003; Sanchez et al., 2012). Using a translational reporter, we confirmed that the procambial pattern of ATHB8 within the SAM was associated with primordium formation following a phyllotactic pattern (Figure 2H). Additionally, we observed that some genes belonging to the Vasculature 1 cluster such *PFA3* and *PFA5* showed an internalized expression pattern converging in primary stem from developing flowers, as observed using the transcriptional reporters *pPFA3::H2B-tdTom* and *pPFA5::H2B-tdTom* (Figure 2I and J). This suggests that unbiased clustering captured the vasculature differentiation process between SAM to primary stem. We observed that *PFA5* and *PFA3* showed the expression patterns expected for vascular bundles characterized during primary growth.

### Overlapping and exclusive role of master regulators MP/ARF5 and DOF5.8 during primordia formation

Several auxin-related genes have been reported to be expressed in early primordium formation. We examined auxin-related genes that were identified as cluster marker genes (Figure 3A). Expression of many of these genes such as *IAA30* and *IAA20* have been previously reported during early primordium formation (Vernoux et al., 2011). *LIKE AUXIN RESISTANT 1* (*LAX1*), an auxin influx carrier gene, has an epidermal and primordial expression pattern observed by *in situ* hybridization and GUS staining in the SAM (Bainbridge et al., 2008). We confirmed its expression at the cellular level using the reporter line *pLAX1::LAX1-YFP* and confirmed the expression of *LAX1* in SAM epidermis, L2 and the EP cluster as seen in our transcriptomic analysis (Figure 3A and 3B). Following the primordial formation pattern, we observed an internalization of LAX1 at the beginning of primordium formation, suggesting the action of LAX1 is to facilitate primordial initiation mainly in very early stages (P1-P2) (Figure 3C). We observed the expression of genes previously characterized during lateral root formation but not associated to early primordia context such as *LATERAL ROOT PRIMORDIA 1* (*LRP1*) (Smith and Fedoroff, 1995), the auxin amido-synthetases *GH3.3* and *YADOKARI 1* (*YDK1/GH3.2*) (Wang et al., 2023). While most auxin-related marker genes are expressed in the EP cluster, some are expressed in different SAM domains such as *ARF6* in the BD (Truskina et al., 2021) and *IAA7* in the Pith cluster, expanding our understanding of auxin-related gene expression beyond early primordium formation.

Contrary to auxin, the expression of genes related to other hormones such as cytokinin (CK) and gibberellic acid (GA) exhibited a dispersed expression across different clusters. Specifically, we noted peak expression of Arabidopsis Response Regulators (ARRs) in distinct cell types: ARR10 in the Vasculature 1 cluster, ARR4 in Vasculature 2, and ARR7 in BD. A similar trend was observed for the LOG gene family, with *LOG1* showing differential expression in Vasculature 1 and *LOG5* in Vasculature 2 (Supplementary Figure 4A). We also observed a heterogeneous expression pattern for GA-related genes such as *GIBBERELLIN 2-OXIDASE 2* (*GA2Ox2*) in the Rib Zone or *GIBBERELLIC ACID METHYLTRANSFERASE 2* (*GAMT2*) in cortex cells (Supplementary Figure 4B).

Given that we captured auxin-related genes co-expressing in a distinct cluster, we investigated the gene regulatory network (GRN) within this group of cells. We considered TF-target interactions that were validated with at least one type of experimental evidence (O’Malley et al., 2016; Alvarez et al., 2019; Brooks et al., 2019) to establish a GRN for the top 100 differentially expressed genes (DEG) in this cluster (Methods). AUXIN RESPONSE FACTOR 5/MONOPTEROS (ARF5/MP) and DOF5.8 were identified as the most connected TFs within this cluster, validating the GRN approach to obtain master regulators for a single cellular identity (Figure 3D) (Supplementary Table II). ARF5/MP has been previously reported as a master regulator of primordium formation in response to auxin and DOF5.8 has been postulated to act downstream of ARF5/MP (Konishi et al., 2015; Larrieu et al., 2022). GO enrichment analysis comparing target genes of ARF5/MP and DOF5.8 showed that they both regulate biological processes related to flower development, pattern specification and regionalization. According to GO analysis, ARF5/MP is primarily involved in processes related to auxin, such as cellular response to auxin stimulus and auxin-activated signalling pathway. In contrast, targets of DOF5.8 are enriched in GO terms such as transcription DNA-templated or regulation of RNA metabolic processes (Figure 3E). The specific role of ARF5/MP in auxin-related functions is further supported by its binding near auxin-related genes, such as INDOLE-3-ACETIC ACID INDUCIBLE 30 (IAA30) and IAA20 (Figure 3F). Moreover, our analysis also highlights the overlapping role of DOF5.8 and ARF5/MP as master regulators of primordial formation through the regulation of a similar set of genes. DOF5.8 and ARF5/MP shared most of the target genes active in early primordium formation such as AHP6 and many others (Figure 3F).

### Stem cortex diverges from vasculature during primordium formation in the SAM

Secondary growth analysis and stained stem sections have shown the presence of cortex in stem tissue, positioned between epidermis and vascular bundles (Sussex et al., 1972; Agusti et al., 2011; Mazur et al., 2016; Shi et al., 2021). However, the precise timing and molecular mechanism governing cortex formation in stems is unstudied. By unbiased clustering, we identified a cluster enriched in the expression of genes that have been previously associated with root cortex such as *JKD*, *AED3*, *HOMEOBOX FROM ARABIDOPSIS THALIANA 7* (*HAT7)* and *C/VIF2* (Figure 4A) (Hassan et al., 2010; Shahan et al., 2022; Nolan et al., 2023). Through analysis of reporter lines *pJKD::JKD-Ypet*, *pAED3::ER-GFP* and *pC/VIF2::H2B-VENUS*, we observed an expression pattern localized to Layer 2 (L2) within the meristematic tissue. Orthogonal views of the SAM using reporter lines revealed cortex-related gene expression during primordium formation (Figure 4B to 4E). Since L1 and L2 layers in the SAM are characterized by anticlinal division (Shapiro et al., 2015; Willis et al., 2016), cortex identity may be generated from L2 cell lineages that originate in the WUS/CLV3 domain. In order to obtain deeper sections of the meristem and primary stem, we assessed *AED3* expression using light sheet microscopy. We observed that the *AED3* promoter is active in a path from early developing primordia downward through the stem (Figure 4B and 4C). The use of light sheet microscopy also allowed us to observe the pattern of *AED3* promoter activation in the rib zone (Figure 4C), as predicted by our clustering analysis (Figure 4A). Thus, we were able to capture a dual expression pattern of *AED3* in the SAM/primary stem region. Gaps in expression in the stem were observed, corresponding to developing flowers we had removed during the dissection process. *JKD* and *C/VIF2* showed a similar expression pattern to *AED3*, with cells expressing the reporter lines in the L2 of flowers and SAM (Figure 4D and 4E).

While *JKD* expression has been previously observed in the SAM (Bahafid et al., 2023), the identification of a group of marker genes with a similar expression pattern confirms a novel domain of cell identity within the SAM that will subsequently generate the cortex in flowers and stem. Following primordium formation in the *JKD* reporter lines, we observed that the expression of this cortex marker gene coincides with morphological changes during primordium formation on the abaxial side of the new organ (Figure 4F). Both EP and Vasculature 1 marker gene expression was also observed during primordium formation (Figure 4F), co-expressing with cortex-related genes within this spatial domain. While vasculature stripes form within the middle of the primordia, cortex cells first appear in Layer 2. Thus, single-cell transcriptomic and cluster validations have allowed us to identify a bifurcation point between two cell identities developing from a joint set within developing primordia.

### S-phase is enriched in floral-related domains

While many molecular players underlying SAM cell cycle dynamics have recently been characterized (Yang et al., 2017; Yang et al., 2021), a comprehensive overall transcriptional description of cell cycle-related gene expression in the SAM remains to be created. We investigated the cell-cycle dynamics of the SAM through trajectory inference (TI) analysis using the two identified cell-cycle related clusters: S-phase and G2/M phase.

We integrated diffusion maps, partition-based graph abstraction and force-directed graphs (Jacomy et al., 2014) (Methods) (Figure 5A and Supplementary Figure 5A). Multiscale diffusion maps were obtained using the Palantir algorithm (Setty et al., 2019), representing differentiation trajectories in pseudotime. We defined an S-phase edge as the initial time-point for the pseudotime analysis (Figure 5A and 5B). Gene expression of histone-related proteins and M-phase genes confirmed that dimensionality reduction captured overall cell-cycle dynamics (Figure 5C). Utilizing cubic spline regression along the cell-cycle trajectory, we identified 245 DEGs with a False Discovery Rate (FDR) < 0.01. Consistent with known cell-cycle dynamics, histone-related genes exhibited an early expression peak in the reconstructed cell-cycle trajectory, succeeded by the expression of G2-associated genes such as *CYCA1;1*, *CDKB2;1* and *CYCB1;1* (Fabian et al., 2000; Menges et al., 2005). Finally, genes associated with M-phase, including *CELL DIVISION CYCLE 20-1* and *20-2* (*CDC20-1* and *CDC20-2*), were expressed in the latter region of the trajectory (Figure 5D and 5E, Supplementary Figure 5B, Supplementary Table III) (Kevei et al., 2011; Yang et al., 2017).

**Figure 5.**
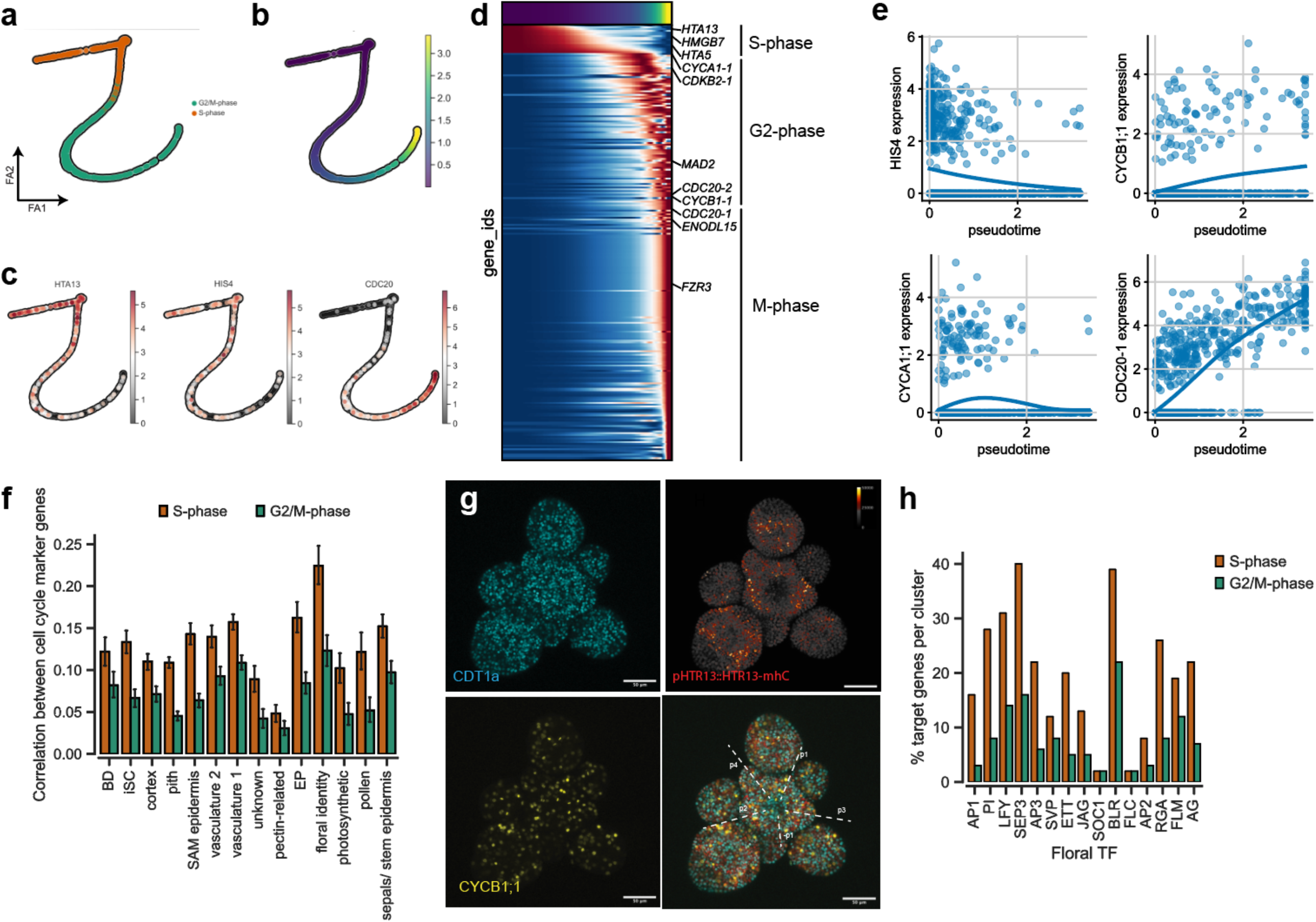
Diverse cell cycle transcriptional profiles in SAM cell types. a) Force-directed graph layout of clusters associated to cell cycle (S- and G2/M-phase). Clusters are represented by different colours. b) Pseudotime estimation of cell-cycle progression. Colour bar represents pseudotime. c) HTA13, HIS4 and CDC20 expression along cell cycle trajectory. d) Heatmap displaying clustering of DEGs based on similarity expression profile along the cell cycle trajectory. e) Expression trends of selected DEGs along the cell cycle progression. f) Bar plot illustrating Pearson correlation between cell cycle marker genes for each cluster and cell cycle clusters. g) Confocal image of SAM from PaClI lines showing CDT1a (G1-phase), HTR13 (S-phase) and CYCB1;1 (G2/M-phase) reporters. S-phase marker line is showed as heatmap of fluorescence intensity. Scale bar, 50 µm. h) Percentage of target genes per cluster for floral homeotic genes based on ChIP-seq datasets. The top 100 expressed genes per cell cycle cluster were assessed using ChIP-seq data.

We then asked whether we could assess the cell-cycle profile of the other identified clusters using the transcriptional profiles of the S-phase and G2/M-phase clusters. Cell-cycle genes expressed in each cell were correlated with the top marker genes from the S-phase and G2/M-phase clusters, separately (Figure 5F) (Methods). We observed distinct correlations between each cluster and the two cell cycle-related clusters, suggesting cell cycle-specific aspects of each cluster. The Pith, Unknown, Pectin-related, Rib zone and pollen clusters showed the lowest correlation with the G2/M-phase cluster, capturing the fact that differentiated cell types undergo fewer cell divisions over time than undifferentiated cells. Unexpectedly, when we compared clusters against the S-phase cluster, we observed that the Floral Identity cluster showed an outlier maximum correlation with S-phase markers gene expression (Figure 5F and Supplementary table IV). Using the PaClI cell cycle reporter line (Desvoyes et al., 2020), we observed that G2/M-phase markers showed a speckled pattern as has previously been reported for a CYCB1;1 translational reporter and other G2/M-related genes (Yang et al., 2017; Yang et al., 2021) (Figure 5G). On the contrary, the histone H3.1 reporter line pHTR13::HTR13-mRFP showed a peak of expression in particular domains such as early primordia and central meristematic floral domains, coinciding with the expression pattern of some floral homeotic proteins (Figure 5G, Supplementary Figure 5C). Thus, although G2/M-phase related genes showed a uniformly speckled expression pattern in the meristem, the expression pattern of some S-phase related genes, though expressed in a speckled pattern, are preferentially distributed in domains associated with the co-expression of floral homeotic genes. Length of S-G2-M-phase has been shown to be flexible depending on the primordium stage (Jones et al., 2017), supporting the heterogeneous expression pattern that we observed for the S-phase reporter line (Figure 5G).

We investigated whether floral homeotic TFs bind to genomic regions of S-phase-expressed mitotic genes. Genome-wide TF binding datasets for most of the floral homeotic genes have been previously reported (Chen et al., 2018). Floral regulator binding sites were found to be highly enriched near S-phase-related genes compared to G2/M-phase-related genes (Figure 5h and supplementary Table V), allowing for the possibility for a direct involvement of floral homeotic factors in regulating S-phase elements. For instance, 58% of the top marker genes in the S-phase cluster are direct targets of SEP3, while only 23% of the G2/M-phase top marker genes are targets of this TF (Figure 5H). We observed similar behaviours for most of the floral homeotic TFs assessed.

### Multiple differentiation events connecting shoot stem cells and primary stem

We then aimed to trace the trajectories of differentiation from stem cells to the various cell types observed in early flowers and primary stem. To increase the resolution of the TI analysis, we removed genes associated with the cell cycle (as they are associated with most of the clusters, and have a trajectory of their own) and re-clustered the cells. Similarly to our previous approach for cell cycle trajectory analysis, we employed diffusion maps, partition-based graph abstraction and force-directed graphs (Methods) to align initial stem cells (iSC, WUS+ cells) with internally located cell types. Based on validated and published expression patterns (Figure 2 and Figure 3), we performed TI analysis between iSC and other identities formed within internal cells of the SAM such as Vasculature 1, Vasculature 2, Cortex and EP (Figure 6A). The number of eigenvectors used to reduce dimensionality were defined when meaningful branches were observed from combined clusters (Supplementary Figure 6). Notably, reducing dimensionality positioned WUS+ cells at one side of the iSC branch (Figure 6B), suggesting the capture of the OC within the trajectory. Additionally, previously identified and validated genes from different cell types were expressed in specific branches of the trajectory, such as *JKD* expressed in cortex branch and *AHP6* expressed in the EP branch (Figure 6C and 6D).

**Figure 6.**
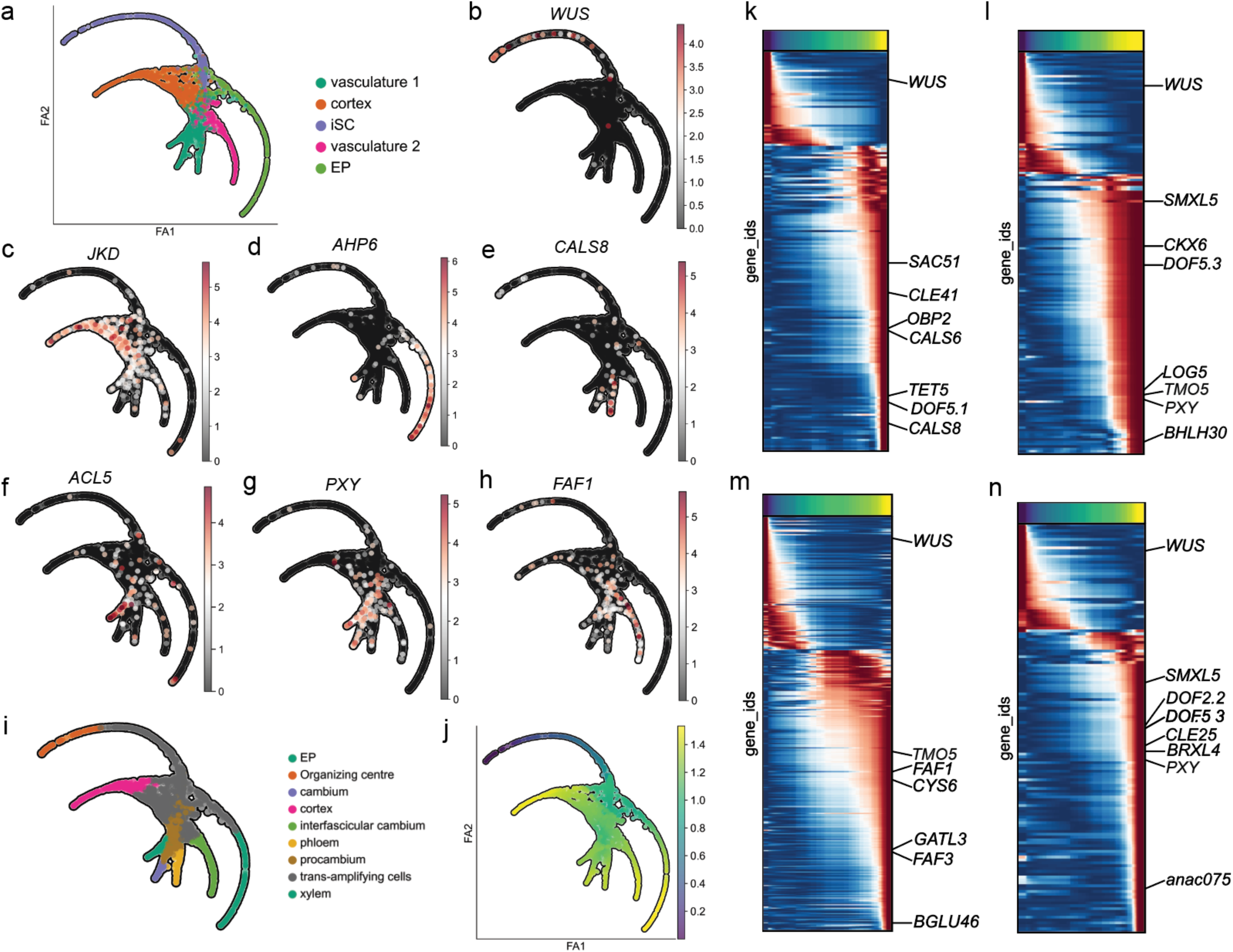
Dynamic transcriptional differentiation of internal cell identities within the SAM and primary stem. a) Force-directed graph layout of clusters associated with internal cell layers such as EP, iSC, Vasculature 1, Vasculature 2 and Cortex. Clusters are represented by different colours b) WUS expression within the force-directed graph layout. c,d,e,f,g,h) JKD, AHP6, CALS8, ACL5, PXY, FAF1 respective expression within the force-directed graph layout. i) Force-directed graph layout of identified cell-types according to branch-specific gene expression. j) Pseudotime analysis using OC-like edge with WUS+ cells as the initial point. k) Heatmap of DEGs along phloem trajectory. l) Heatmap of DEGs along xylem trajectory. m) Heatmap of DEGs along interfascicular cambium trajectory. n) Heatmap of DEGs along cambium trajectory Heatmap of DEGs along xylem trajectory

TI analysis separated the Vasculature 1 cluster into three different branches with distinct transcriptional profiles. One branch showed the expression of genes whose functions are related to development of phloem such as *CALLOSE SYNTHASE 8* (*CALS8*) (Figure 6E) (Ross-Elliott et al., 2017), another branch showed the expression of genes related to development of xylem vessels such as *ACAULIS 5* (*ACL5*) (Figure 6F) (Muñiz et al., 2008), and a third branch showed expression of genes whose transcription is associated with the cambium, such as *PHLOEM INTERCALATED WITH XYLEM* (*PXY*) (Figure 6G) (Kucukoglu et al., 2017). The Vasculature 2 cluster did not split into branches and it was defined by the expression of vascular genes such as *FAF1* and *FAF3*, both having an internalized expression pattern not related to phyllotaxis that suggest the identification of interfascicular cambium (Wahl et al., 2010) (Figure 6H). Thus, trajectory analysis enabled us to reconstruct the spatiotemporal connection between cell located in the OC (WUS+ cells) and differentiated cell types observed in primary stem such as cortex, xylem, phloem, cambium and interfascicular cambium (Figure 6I).

To identify differentially expressed genes along a particular trajectory, we conducted a pseudotime analysis, defining WUS+ cells as the starting point (Figure 6J). Gene expression was modelled as a function of pseudotime and a cubic spline regression was used for each branch independently to identify genes whose expression was significantly associated with the trajectory (Methods). Using a false discovery rate (FDR) < 0.01, we identified 190 DEGs during phloem differentiation (Figure 6K), 171 DEGs along xylem differentiation (Figure 6I), 368 DEGs for interfascicular cambium (Figure 6M), 143 DEGs along the fascicular cambium trajectory (Figure 6N), 385 DEGs in cortex differentiation (Supplementary Figure 7A) and 33 DEGs along EP differentiation (Supplementary Figure 7B and Supplementary table VI).

Cortex-specific gene expression, including from *JKD*, *C/VIF2* and *AED3*, was found early and expression was maintained throughout the cortex branch (Supplementary Figure 7A). The vasculature 2 or interfascicular cambium trajectory exhibited a similar expression pattern with early-expressed interfascicular RNAs such as the transcripts of *FAF1* and *FAF3* and of late-expressed interfascicular genes such as *BETA GLUCOSIDASE 46* (*BGLU46*), the product of which participates in lignification, a process observed in interfascicular vasculature (Figure 6M) (Escamilla-Treviño et al., 2006). Within the phloem trajectory we observed the expression of genes characteristic of differentiated phloem such as *CALS8*, *CALS6* (Ross-Elliott et al., 2017), *OBP2* (Skirycz et al., 2006), *CLE41* (Etchells and Turner, 2010) and *PEAR2/DOF5.1* (Miyashima et al., 2019) (Figure 6K). The xylem-related trajectory showed *TMO5*, *TMO5L* (De Rybel et al., 2013), and *PXY* (Etchells and Turner, 2010), as differentially expressed genes (Figure 6I). The cambium trajectory also manifested upregulation of *PXY* but it did not show the activation of xylogenesis-expressed genes such as *TMO5* (Agusti et al., 2011) (Figure 6N). We observed that some genes were expressed before any bifurcation point such as *TMO5* and *ATHB8*, defining a procambial region within the trajectory that has been previously observed (Figure 2 and Supplementary Figure 7C) (Mor et al., 2022). Thus, we were able to capture the transcriptional landscape of differentiation processes for a variety of cell identities defined between the SAM and primary stem.

## DISCUSSION

Leveraging single-nucleus transcriptomics, we provide a quantitative characterization of cell types and differentiation processes occurring within the shoot apical meristem and primary stem in *Arabidopsis thaliana*. By extracting gene regulatory networks we were able to capture the gene expression dynamics underlying floral primordium formation in considerable detail. Moreover, we decoded the dynamics of the cell cycle, revealing spatially heterogeneous features linking to S-phase cells and floral homeotic genes. Combining internally-located cell identities, we achieved a detailed description of the differentiation process from shoot stem cells through several cell identities observed within primary stem such as cortex, phloem, xylem, cambium and interfascicular cambium (Figure 7).

**Figure 7.**
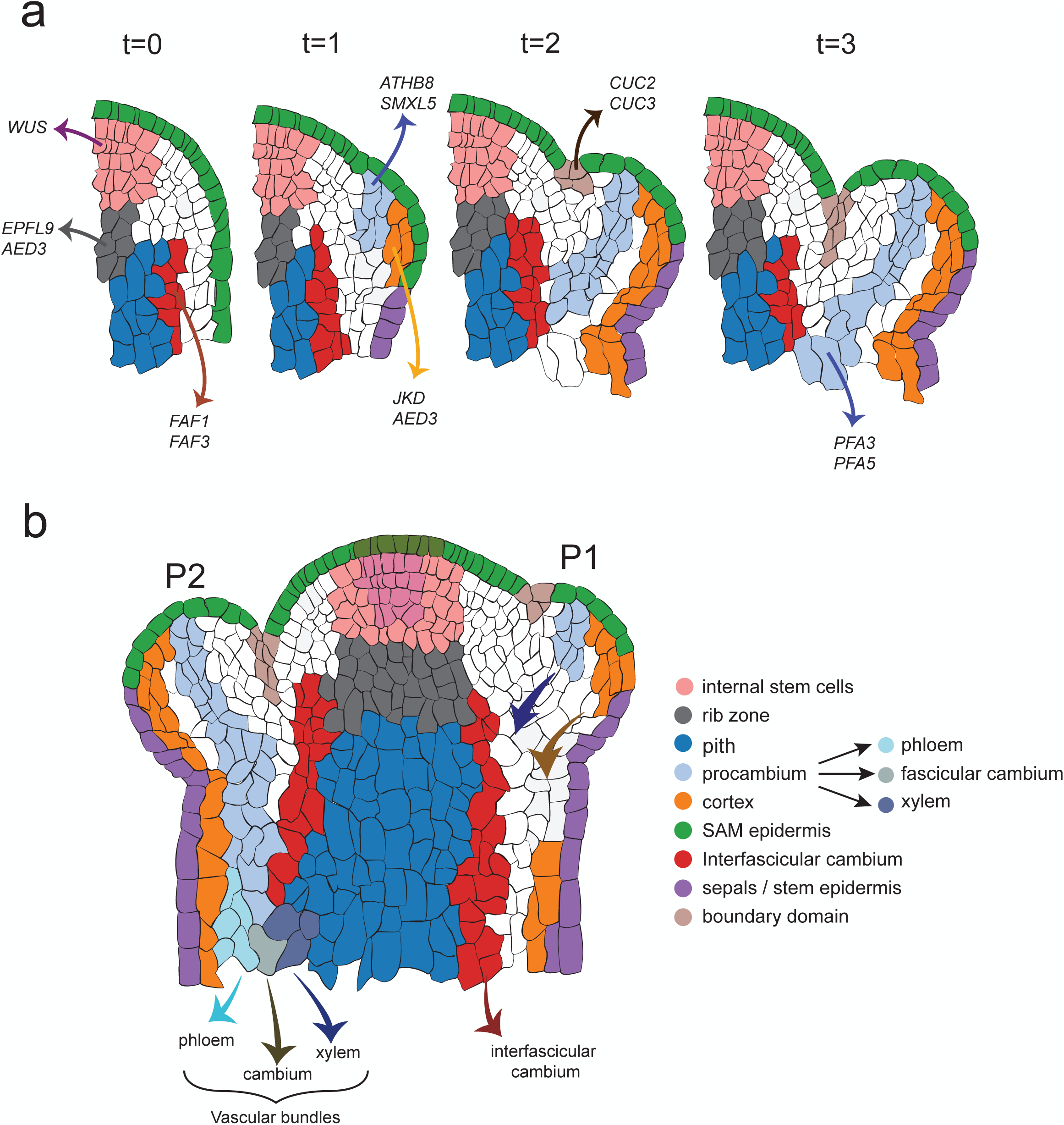
Cell identities and differentiation trajectories within the SAM and primary stem. a) Dynamics of cell types during primordia formation. Identified marker genes per domain are labelled, and different cell-types are represented by different colours. Procambial strands and cortex-expressed genes are highlighted during primordia formation. b) Primary stem and SAM cell identities. Procambial strand observed in developing primordia form vascular bundles, which differentiate into phloem, cambium and cortex within the primary stem.

### Unbiased clustering captured SAM cell heterogeneity with unprecedented precision

Previous single-cell RNA-seq experiments have characterized the apex of *Arabidopsis thaliana* (Zhang et al., 2021a; Xu et al., 2024). However, the differentiation processes from stem cells to the heterogeneity of cell types within the apex have remained uncharted. Our work underscores the utility of dissecting small tissue regions to capture the stem cell population for sequencing, allowing extraction of the subsequent differentiation processes in the SAM of wild type plants. We identified 16 different cell identities with characteristic transcriptional profiles within the SAM and primary stem. Unbiased clustering did not identify a distinct peripheral zone as has been proposed (Meyerowitz,1997; Murray et al., 2012), suggesting that once cells have exited the OC region, they rapidly undergo cell differentiation towards the different cell types observed in primary stem such as vasculature or cortex (Figure 2 and Figure 4). Alternatively, increasing the resolution by adding more nuclei to our analysis could help us to increase the variety of cell identities observed in the SAM.

We expanded the repertoire of auxin-related genes within the early primordia cluster and validated that previously uncharacterized marker genes such as the influx carrier *LAX1*, expressed in L1 and L2 in developing primordia. Although *lax1*-related mutants can generate new primordia, the phyllotaxis pattern is disrupted (Bainbridge et al., 2008), supporting its role in generating auxin maxima at the correct location (Heisler and Jönsson, 2007). Moreover, the detection of marker genes mostly expressed in each cluster allowed us to decode the gene regulatory network behind primordium formation, proposing ARF5/MP and DOF5.8 as the main regulatory factors in control organ formation (Figure 3). Mutant lines of DOF5.8 do not show apparent phenotypes, but they enhance the phenotypes of the weak alleles of other genes, such as *arf5-2* or *mp^S319^* (Konishi et al., 2015; Larrieu et al., 2022). Despite sharing several target genes, the distinct phenotypes observed in DOF5.8 and ARF5/MP mutant backgrounds can be attributed to the broader auxin-related target genes of ARF5/MP, like *IAA30* and *IAA20*.

Validation of several reporter lines revealed that the previously unidentified cortex cell identity is generated in the L2/L3 during primordium formation (Figure 4). *JKD* expression has been previously observed within this domain of meristematic tissue (Bahafid et al., 2023). We confirmed that *JKD* expression pattern as indicative of a new cell type by identifying additional cortex-expressed genes with similar expression pattern such as *AED3* and *C/VIF2*. SAM size in *jkd-4* mutant lines is larger compared to WT due to greater cell proliferation, suggesting a role of stem cortex in stabilizing SAM size (Bahafid et al., 2023). Further cellular and mechanical characterization of cortex-related mutants could be relevant to understanding the role of this cell layer within the above-ground tissues of flowers and stem.

### Spatiotemporal control of S-phase and cell fate commitments in the SAM

Clusters enriched in cell cycle gene expression reveal significant transcriptional changes during cell cycle phases, enabling their categorization into distinct clusters. Correlating cell cycle marker genes showed that SAM cell types exhibit specific cell cycle dynamics, similar to what has been observed in other organisms and tissues (Pauklin and Vallier, 2013; Dalton, 2015). Expression of S-phase reporter lines are primary seen in floral identity domains, with a higher correlation between S-phase marker genes and the Floral Identity cluster (Figure 5G), supported by a higher enrichment for S-phase marker genes as target of floral homeotic genes according to ChIP-seq datasets (Figure 5H) (Chen et al., 2018). While the functional impact of heterogenous S-phase in different SAM domains remains unknown, S-phase regulation could influences morphogenesis and cell fate decisions, as seen in various aspects of animal development such as development of *Drosophila* central nervous system (Weigmann and Lehner, 1995), regulation of cell fate switches during erythroid differentiation (Hwang et al., 2017) and between neural progenitors in developing neocortex in mammals (Soufi and Dalton, 2016; Turrero García et al., 2016). Understanding whether the S-phase enriched regions in floral domains are formed due to shorter M-phase or prolonged S-phases could shed light on the role of S-phase in shaping plant morphology.

Utilizing WUS+ cells as a proxy for undifferentiated cells, we delineate the differentiation transcriptional profiles for various cell identities such as cortex and vasculature (Figure 6). In comparison to the to the root apical meristem (RAM), where cells achieve their terminal identity after a few divisions upon exiting the quiescent centre (Pardal and Heidstra, 2021), our observations confirmed that cells in the SAM must attain specific locations to assume a definitive cell fate, such as procambium strands or cortex within the developing primordia. Cell fates in plants are determined by cell lineage inheritance and positional information (van den Berg et al., 1995; Yu et al., 2017). Unfortunately, our trajectory analysis could not differentiate between the options of cell fate commitment. The use of clone lineage tracking analysis using the repertoire of identified marker genes per cluster could help us to decode the mechanism underlying cell fate commitment in above-ground organs. Nevertheless, by elucidating the differentiation transcriptional profiles within the SAM, our study characterizes several cell fate differentiation processes in shoot meristematic tissue (Figure 7), paving the way for future investigations into the dynamic nature of cell identity and development in plants.

## MATERIAL AND METHODS

### Plant material and growth conditions

Seeds were sterilized using 70% (v/v) alcohol for 7 minutes and subsequently plated on 1/2 Murashige and Skoog basal medium (MS) (pH = 5.7). Seeds were stratified in the plates for 48 hours at 4°C in darkness and plates were then positioned horizontally in a growth chamber set to 22°C with a photoperiod of 16 hours light and 8 hours dark. Ten-day-old seedlings were transplanted into 9 cm pots filled with Levington F2 compost and placed in a growth chamber set to 22°C, 16 hours light/8 hours dark.

### Nuclear isolation and 10X snRNA-seq library preparation

A total of 750 dissected Col-0 shoot apical meristems from three biological replicates (100, 350 and 300 meristems respectively) were used for the single-nucleus RNA-seq approach. Meristems were dissected shortly after bolting (with stem length of 1-3 cm and flowers beyond stage 3 were removed). The stem was cut just below the SAM (∼ 150-200 µm, Fig. 1A), and tissues were collected and frozen in an Eppendorf tube using liquid nitrogen. For each sample batch, 400 µl of Honda buffer (2.5% Ficoll 400, 5% Dextran T40, 0.4M Sucrose 10 mM MgCl_2_, 1 µM DTT, 0.5% Triton X-100, 1 tablet/50 ml cOmplete Protease Inhibitor Cocktail (Roche), 0.4 U/µl Ribolock (Thermo Fisher Scientific), 20 mM Tris-HCl, pH 7.4) were added, and the tissue was mechanically disrupted using a motorized pestle for 10 seconds to release the nuclei. Subsequently, 500 µl of Honda buffer was added to the tubes, and the resulting suspension was filtered through a 70-µm strainer (Sigma) and then filtered again through a 40-µm strainer (Sigma). The filtered solution was centrifuged at 1500 g for 6 min at 4°C. The pellet was resuspended carefully in 300 µl of landing buffer (1X PBS with 4% BSA, 2U/µl Ambion RNase Inhibitor, 1U/µl Superasein RNase inhibitor). Nuclei were stained with propidium iodide (PI) and sorted by gating on PI peak intensity using a BD Bioscience Influx cell sorter with a 100 µm nozzle at 20 psi pressure. The PI signal was detected with a 75 mW 561 nm laser using a 670/30 bandpass filter. 10,000 - 40,000 nuclei per sample were collected in 200 µl of landing buffer and centrifugated at 1500 g for 6 min at 4°C. The nuclear pellet was resuspended in 43 µl of landing buffer and loaded to a 10X Chromium chip for library preparation using Chromium Single Cell 3’ Reagent Kits v3 according to the manufacturer’s instructions. Library were sequenced using Illumina NovaSeq 6000 Sequencing System.

### snRNA-seq data analysis

Fastq files were processed with Cell Ranger v3.1.0 (Zheng et al., 2017), using default parameter values and reads were aligned to the *Arabidopsis thaliana* TAIR10 reference genome (https://www.arabidopsis.org/download/list?dir=Genes%2FTAIR10_genome_release). Ambient mRNA (nucleus-free RNA molecules in the cell suspension) was removed using the CellBlender software package (https://github.com/broadinstitute/CellBender) (Fleming et al., 2023). Filtered genes-by-cell matrices for each batch were concatenated and analysed using Scanpy package (Wolf et al., 2018). To identify doublets, we applied Scrublet with parameters: expected_doublet_rate = 0.05, min_counts = 2, min_cells = 3, min_gene_variability_pct =85 and n_prin_comps = 30 (Wolock et al., 2019). Cells were filtered to remove the 1% bottom and top outliers in terms of minimum genes, maximum genes, maximum reads and minimum reads. Additionally, 1% outlier genes with maximum counts were removed. Gene transcripts present in fewer than 5 cells and organellar genes were excluded for further analysis. We normalized the datasets for cell library size to 10,000 counts without excluding highly expressed genes, log-transformed and selected 1000 top HVGs using the ’seurat_v3’ algorithm for the dispersion-based method. We regressed out the total counts per cell as a source of variation, and gene counts were scaled to mean zero and unit variance. For data visualization, we selected 50 principal components with 100 neighbours. Batch samples were integrated with bbknn (Polański et al., 2020), and we performed a uniform manifold approximation and projection (UMAP) using Leiden community detection with a 0.9 resolution (Supplementary Table I). A Wilcoxon rank sum test with tie correction and Benjamini-Hochberg p-value correction method was conducted to identify marker gene transcripts (Pullin and McCarthy, 2024).

#### Gene ontology (GO)

We used the ShinyGO 0.80 platform (http://bioinformatics.sdstate.edu/go/) for GO analysis. ShinyGO calculated false discovery rate based (FDR) on nominal p-value from hypergeometric test. The top 100 marker gene transcripts per cluster were used for GO analysis.

#### Comparison with poplar spatial transcriptome

Each gene detected in the poplar spatial transcriptomics from shoot apex and stem was associated to a particular cluster based on its peak of expression (Du et al., 2023). Arabidopsis orthologs were defined by Du and collaborators. Subsequently, top marker genes from our datasets were compared with cluster-associated genes from the spatial transcriptomics dataset.

#### Early primordia gene regulatory network

*S*ince the top 100 marker genes within a cluster share gene expression pattern within the same group of cells, we cross-referenced the top 150 marker genes with available TF-binding information from DAP-seq, ChIP-seq and TF-TARGET (Supplementary Table II) (O’Malley et al., 2016; Alvarez et al., 2019; Brooks et al., 2019).

### Trajectory analysis

To infer developmental trajectories in clusters of interest, data were extracted from the integrated Scanpy objects and analysed using the Palantir (Setty et al., 2019) and scFates packages (Soldatov et al., 2019; Faure et al., 2020; Faure et al., 2023). To linearly reduce the dimensionality of clusters, we first obtained the diffusion component using Palantir ’run_diffusion_maps’ function from the 50 principal components obtained for the cluster of interest. Subsequently, a multiscale-data matrix was generated using defined number of eigen vectors from the diffusion map using ’determine_multiscale_function’ from Palantir. Three and nine eigen vectors were used for cell cycle and internal cell types trajectories respectively. Using the obtained multiscale-data matrix, we computed the neighbourhood graph for each cell using scanpy.pp.neighbors function with 30 nearest neighbours. Then, a force-directed graph using the first two principal components was employed to infer trajectories with default parameters (Jacomy et al., 2014).

Along the obtained trajectory, a principal tree was inferred using scfates.tree function (method = ’simple ppt algorithm’) (Soldatov et al., 2019). Each cell is assigned a probability of belonging to different nodes in the tree. Each node of the tree contains values for all cells, which each cell having an assignment strength to a node between 0 and 1, where 1 indicates the closest proximity to the node. Root cells were selected depending on the trajectory topology (S-phase or WUS+ cells for cell-cycle and internal cells trajectories respectively). Pseudotime values were generated as a distance on the tree from the selected roots and projected to cells.

To identify the set of genes significantly associated with the trajectory, we employed the ’scfates.tl.test association’ function from the scFates package. This function models feature expression as a function of pseudotime for each branch independently using a cubic spline repression (*g_i_ ∼ t_i_)*. This tree-dependent model is then compared to an unconstrained model (*g_i_ ∼ 1*), where gene expression is assumed to be independent of pseudotime, using an F-test to evaluate the significance of the association. Genes with significant associations were then fitted using a generalized additive model (GAM) to derive smoothed expression trends over pseudotime (scFates.tl.fit). These fitted trends were visualized as a heatmap (’scfates.pl.trends’), ordered by maximum fitted value of each feature along the pseudotime trajectory (Supplementary Table III and VI).

#### Cell-cycle transcriptional profile per cluster

For each cell cycle cluster, we calculated the average expression level of the top 100 marker genes within their respective clusters. For each individual cell, we computed the Pearson correlation between its expression profile (using the top 100 marker genes) and the average expression profile of the marker genes within the corresponding cell cycle cluster (Supplementary Table IV).

### Microscopy analysis of fluorescent markers

#### Confocal microscopy

Shortly after bolting (stem length ∼ 1cm), the shoot apices were cut and SAMs were dissected removing most of the flowers as it was previously described (Yang et al., 2017). The apices were transferred to a small petri dish containing MS medium (1/2 Murashige and Skoog basal media) (pH = 5.7). Once in the plates, the SAMs were submerged in water and the remaining flowers were removed. To show cell boundaries, each SAM cell wall was stained with 10 µl of Propidium Iodide (PI) 1 mg/ml solution (w/v) for 10 minutes. Confocal z-stacks of SAMs were acquired with Zeiss LSM880 Confocal microscope using a 25 x NA 0.95 water dipping objective. Laser excitations were 488 nm (GFP, YFP) and 561 nm (RFP,PI). Fluorescence intensity was measured in Fiji Image J.

#### Light-sheet microscopy

Light-sheet imaging was performed on a custom-built light sheet microscope (Rowe et al., 2023). Excitation and emission water immersion objectives are arranged on a horizontal plane, perpendicular to each other so that their focal volumes coincide. The sample is mounted upside down in a pipette tip filled with agarose with the part to be imaged is protruding at the bottom. The excitation objectives (Nikon 10x CFI Plan Fluorite Objective, 0.3 NA) and the 2 emission objectives (Olympus 20X XLUMPLFLN 20x, 1.0 NA) are facing each other respectively. The light sheet microscope is a galvanometer scanned light sheet microscope with a vertically scanned gaussian laser beam. The images are recorded with a Hamamatsu Orca Flash 4 camera (Hamamatsu Orca Flash4 V2) with bandpass filter placed in front of Chroma ET525/50m. to detect only the fluorescence. Camera exposure time was set to 100ms per frame and the excitation laser excitation power was set to @488nm for GFP. Stacks of 400 planes with a step size of 1um were typically recorded.

## ACKNOWLEDGEMENTS

We express our gratitude to Dr. Patrick Laufs for the pSOD7::GFP reporter line, Dr. Pauline E. Jullien for the pAGO9::mCherry-AGO9 reporter lines, Dr. Tatsuo Kakimoto for the pPFA5::H2B--Tom and pPFA3::H2V-tdTom lines, Dr. Donghwi Ko for assistance with the pAHP6::ER-GFP, Dr. Pawel Roszak for sharing the pATHB8::ATHB8-GFP, Dr. Danjan Swarup for sharing pLAX1::LAX1-VENUS reporter line and Dr. Philip Benfey for providing pC/VIF2::H2V-Venus, pAED3::ER-GFP (pCORTEX) and pJKD::JKD-Ypet reporter lines. We also thank Louis Faure for comments and feedbacks on the use of scFates package. We thank Dr. Reiner Shulte and Dr. Gabriela Grondys-Kotarba for their great assistance using Flow Cytometry. We thank CRUK Cambridge Institute and specially Dr. Wing-Kit Leung and Dr. Katarzyna Kania for conducting 10X libraries and Illumina sequencing. We thank Dr. Ray Wightman and Mr. Gareth Evans for constant assistance with confocal imaging. Finally, we also would like to thank the entire professional service community at SLCU for making this work possible.

## AUTHOR CONTRIBUTIONS

Conception and Design of Experiments: SM, HJ, EM, JL.

Confocal Imaging: SM

Light-Sheet imaging: ML, SM

Interpretation of results: SM, HJ, EM, JL

Writing Manuscript: SM

Editing Manuscript: SM, HJ, EM, JL

