## Supplementary Figures for "Single-nucleus transcriptomics resolves differentiation dynamics between shoot stem cells and primary stem"

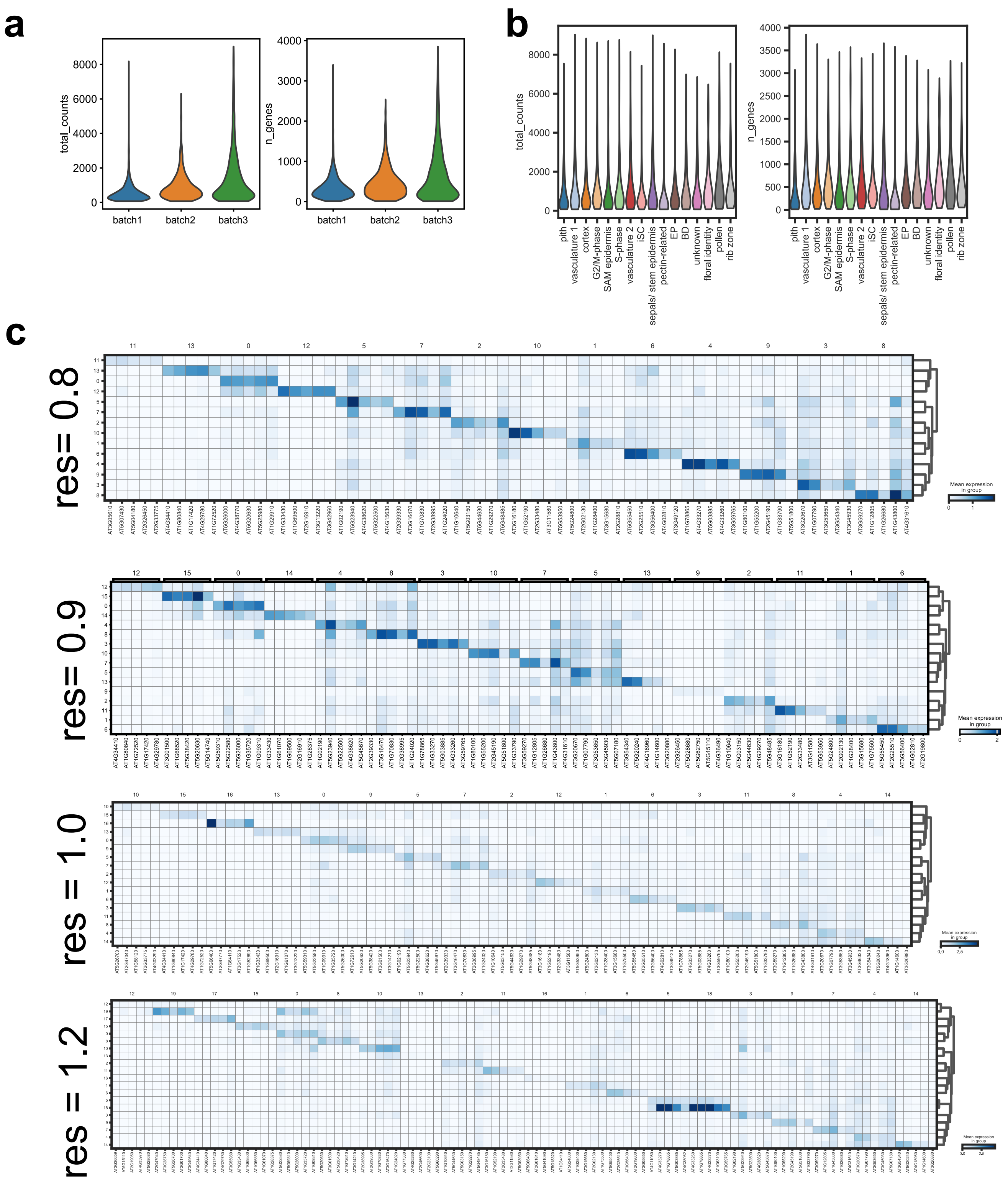


**Supplementary Figure 1. Summary of single-nuclei RNA-seq batches from SAM.** a) Violin plot displaying the total number of reads and genes per batch. b) Violin plot illustrating the total number of reads and genes per cluster after integrating three batch samples. c) Matrix plot highlighting the top 5 marker genes per cluster, identified using the Wilcoxon test, at various cluster resolutions. A resolution of 0.8 was selected for subsequent analysis.

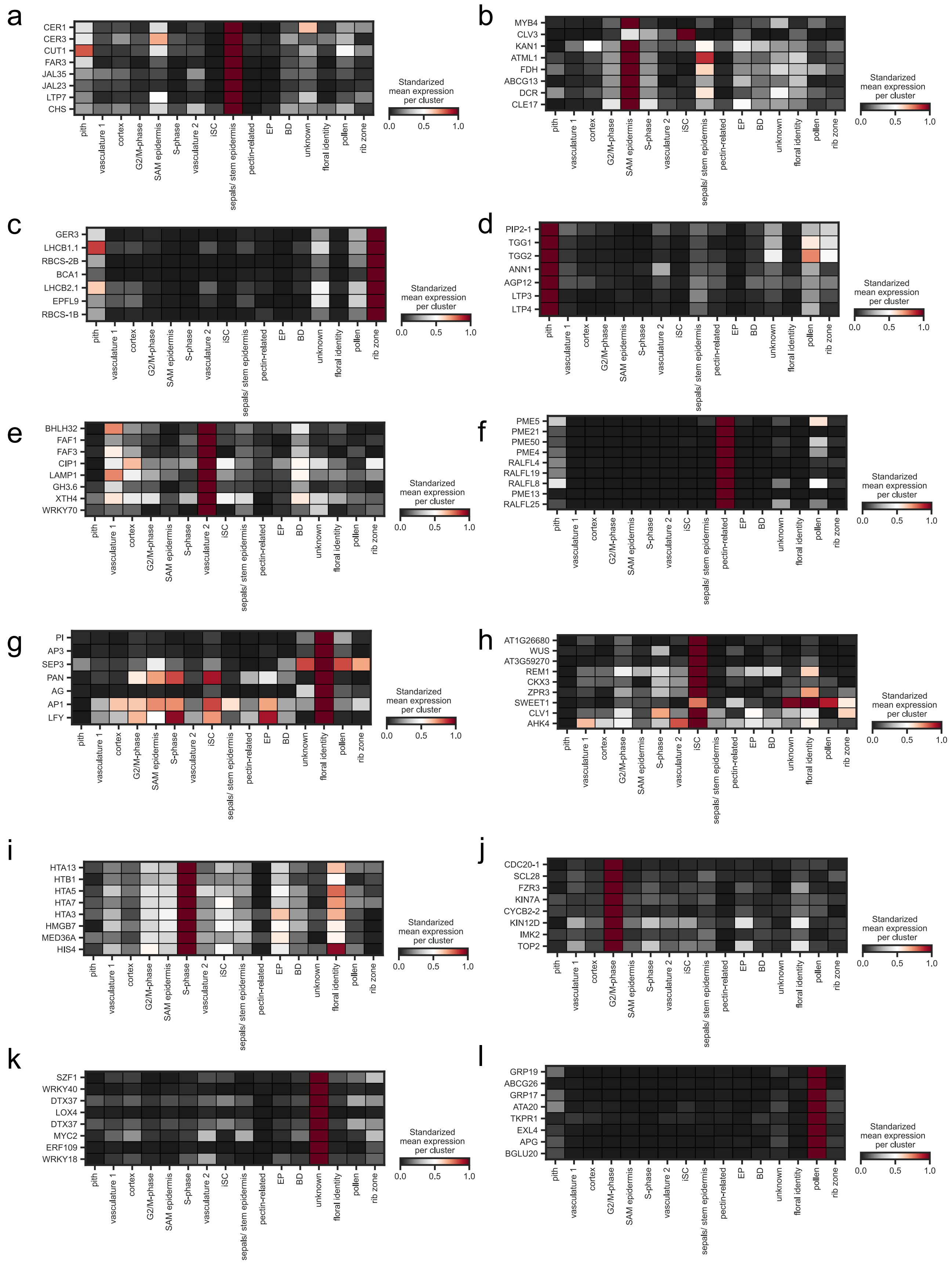

**Supplementary Figure 2. Differentially expressed genes per clusters.** Heatmap plots illustrating selected marker genes from different clusters. Each gene is individually normalized subtracting its minimum value from all values and then dividing each value by its maximum value. a) Stem/Flower epidermis cluster. b) SAM epidermis cluster c) Rib zone cluster. d) Pith cluster. e) Vasculature 2 cluster. f) Pectin-related cluster. g) Floral identity cluster. h) iSC cluster. i) S-phase cluster. j) G2/M-phase cluster. k) Unknown cluster. i) Pollen cluster.

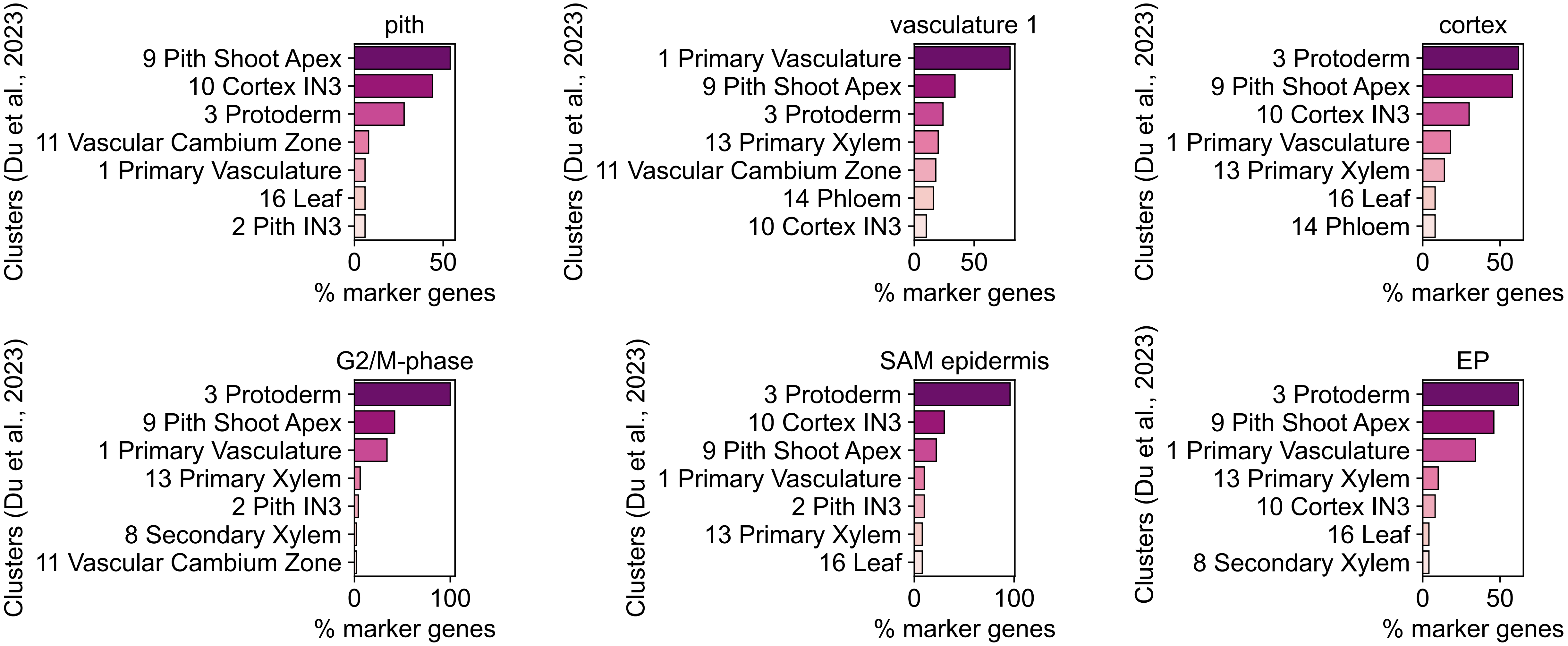


**Supplementary Figure 3. Comparison with available spatial transcriptomics from the poplar shoot apex.** Top 100 marker genes from the SAM atlas clusters were compared to genes expressed in spatial transcriptomics by Du et al., 2023. Genes were associated to each cluster in the spatial transcriptomic data based on their peak expression levels. Subsequently, the marker genes from the SAM atlas were compared to the genes with the associated cluster from spatial transcriptomics. Ortholog recognition between poplar and Arabidopsis was performed by Du and collaborators.

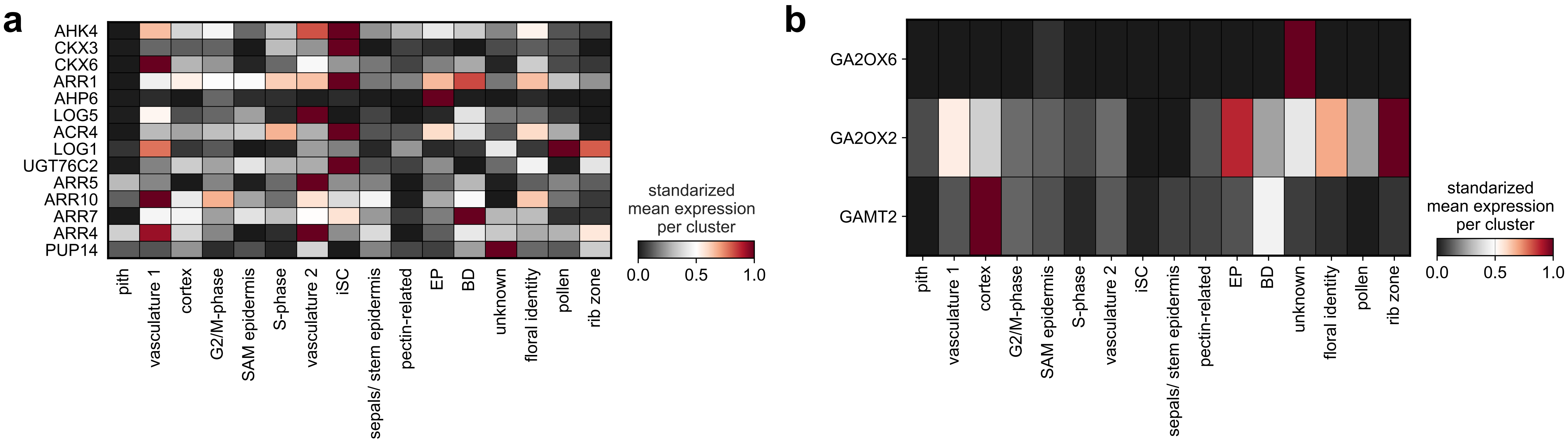


**Supplementary Figure 4. Expression pattern of cytokinin and Gibberellic acid (GA)-related genes.** a) Heatmap plot illustrating normalized expression values of cytokinin-related marker genes. b) Heatmap plot illustrating normalized expression values of GA-related marker genes.

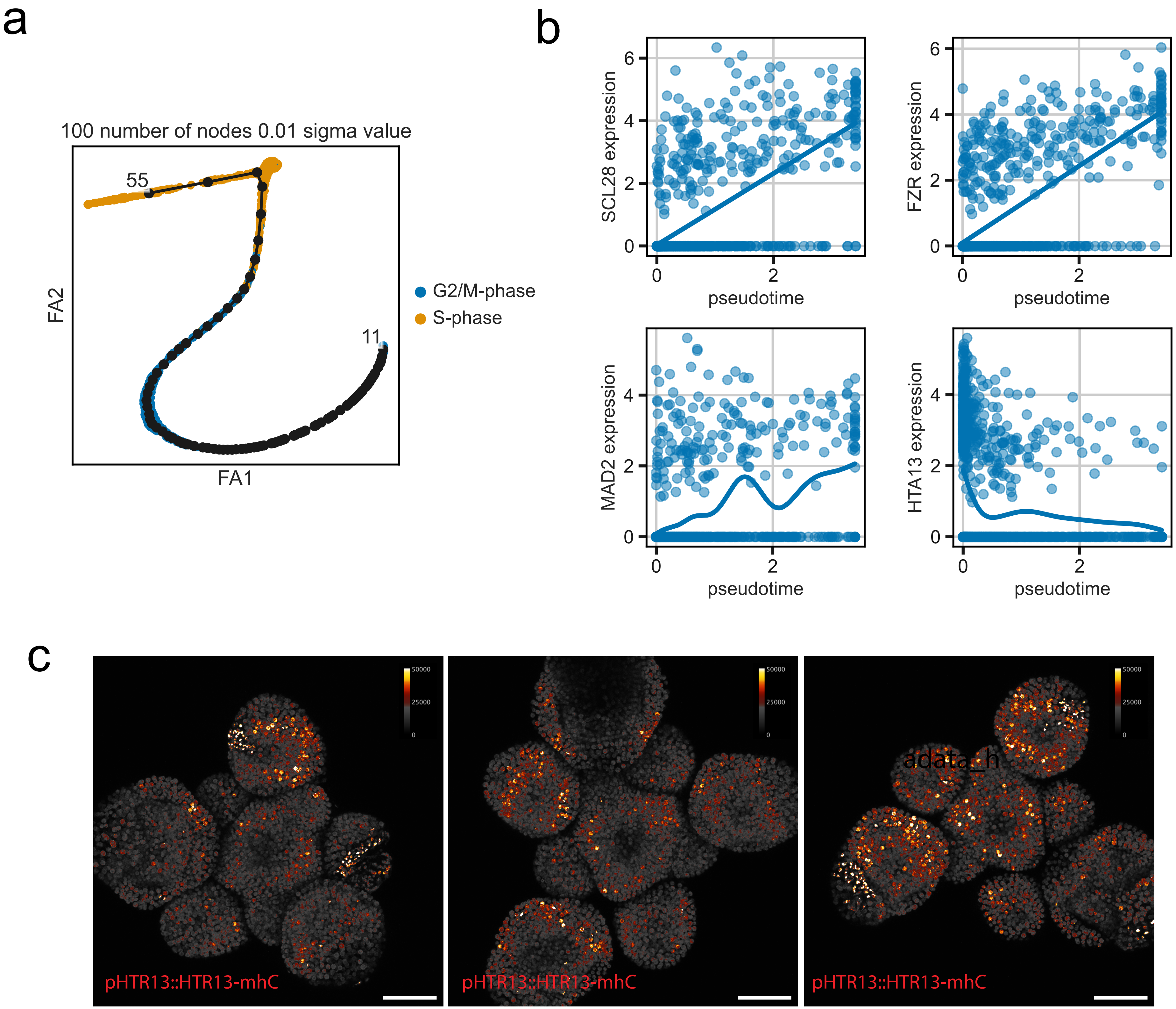


**Supplementary Figure 5. Dynamics of differential gene expression along the SAM cell cycle.** a) Force-directed graph layout depicting nodes and sigma value used to cover the trajectory according to scFates analysis (Methods). b) Gene expression trends of differentially expressed genes along the cell cycle trajectory. Scatterplots indicate gene expression per cell, and expression trends are depicted with fitted expression levels. c) Maximal projection of SAM from three PaCII lines displaying the intensity of fluorescence in the S-phase reporter line.

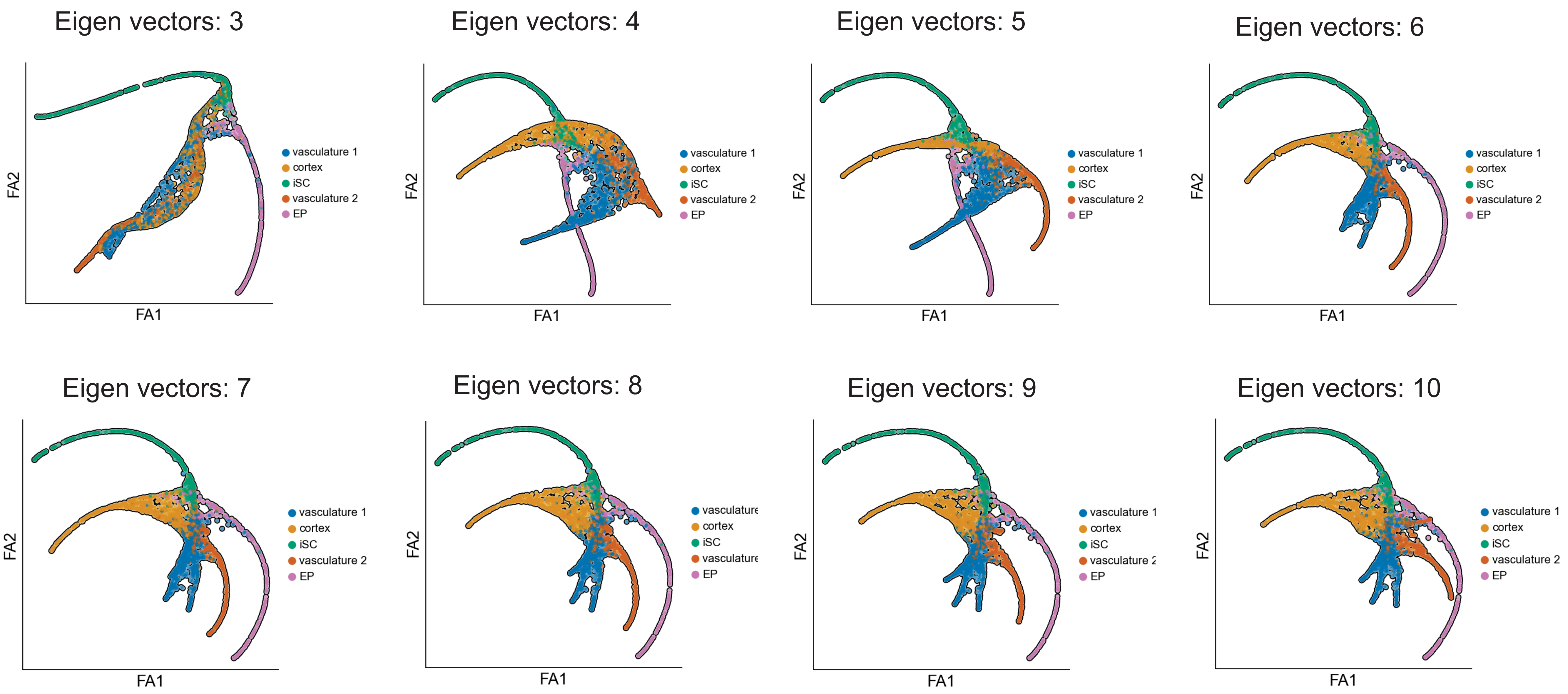


**Supplementary Figure 6 Dimensionality reduction analysis for internal cell clusters.** Force-directed graph layout of clusters associated to internal cell layers such as EP, iSC, Vasculature 1, Vasculature 2 and Cortex using different eigenvectors. An eigenvector of 9 was used for the analysis shown in Figure 6 and described in the text.

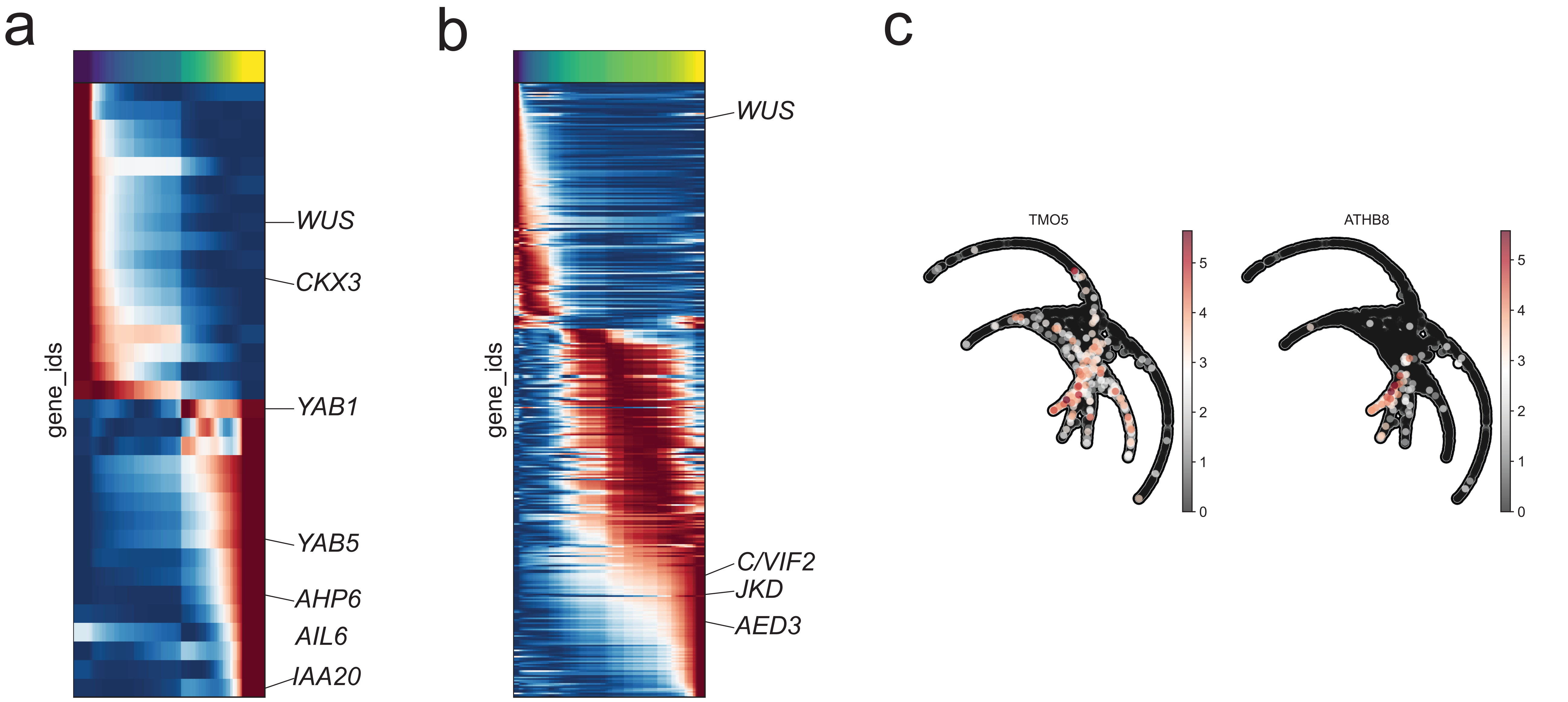


**Supplementary Figure 7. Differentiation trajectory of EP and cortex clusters.** a) Heatmap of DEGs along early primordia differentiation. b) Heatmap of DEGs along cortex trajectory. Some cortex-expressed genes are highlighted. c) Heatmap labelling the expression per cell of *CLE41*, *PIN1*, *TMO5* and *ATHB8* in the force-directed graph layout trajectory.
